# Inner membrane components of the plasmid pKM101 type IV secretion system TraE and TraD are DNA-binding proteins

**DOI:** 10.1101/2024.08.18.608465

**Authors:** Zakaria Jemouai, Aleksandr Sverzhinsky, Jurgen Sygusch, John Pascal, Christian Baron

## Abstract

The increase of antimicrobial resistance constitutes a significant threat to human health. One of the mechanisms responsible for the spread of resistance to antimicrobials is the transfer of plasmids between bacteria by conjugation. This process is mediated by type IV secretion systems (T4SS) and previous studies have provided *in vivo* evidence for interactions between DNA and components of the T4SS. Here, we purified TraD and TraE, two inner membrane proteins from the *Escherichia coli* pKM101 T4SS. Using electrophoretic mobility shift assays and fluorescence polarization we showed that the purified proteins both bind single-stranded and double-stranded DNA in the nanomolar affinity range. The previously identified conjugation inhibitor BAR-072 inhibits TraE DNA binding *in vitro*, providing evidence for its mechanism of action. Site-directed mutagenesis identified conserved amino acids that are required for conjugation that may be targets for the development of more potent conjugation inhibitors.

## Introduction

Conjugation is a mechanism of horizontal gene transfer between bacteria, and it requires the presence of type IV secretion systems (T4SS) [1]. These are large nanomachineries that span the entire cell envelope, allowing biogenesis and formation of pili that mediate contact with recipient cells, followed by DNA transfer. T4SS are coded on self-transmissible conjugative plasmids that contain genetic elements for the T4SS machinery (minimal T4SS) and the DNA processing machinery (relaxosome complex).

The minimal T4SS comprises 12 proteins (VirB1 to VirB11 and VirD4), named after the components of the *Agrobacterium tumefaciens* T4SS [2]. The outer membrane complex is composed of VirB7, VirB9 and VirB10 and the inner membrane and periplasmic complex is composed of VirB3, VirB5, VirB6 and VirB8. The machinery is powered by three ATPases: VirB4, VirB11 and VirD4 that associate with the inner membrane. A recently published cryo-electron microscopy (cryo-EM) structure revealed the subatomic architecture of the R388 T4SS and confirmed interactions between proteins that had previously been shown using various biochemical techniques ([3], [4], [5], [6]). The extracellular pili are formed by VirB2 and VirB5 and they also contain phospholipids in their structure [7–11]. Recent work on the *A. tumefaciens* and *E. coli* P-type pilus structure [8, 9] revealed the presence and importance of positive charges in the lumen. This observation raised the question whether DNA is transferred through the pilus. Indeed, recent work provided evidence for this notion using an F-type pilus [12, 13].

There is also indirect evidence for the binding of the translocated DNA to components of the T4SS. Using chemical crosslinking and immunoprecipitation of transferred DNA (TrIP) it was shown that VirB2, VirB6, VirB8, VirB9 and VirB11 may bind DNA, indicating a possible DNA transfer pathway through the *A. tumefaciens* T4SS [14].

Here, we directly tested this hypothesis using purified homologs of VirB6 and VirB8 from the plasmid pKM101 T4SS. In our previous work we showed by negative stain EM and small-angle X-ray scattering (SAXS) that the inner membrane protein VirB8 homolog TraE forms hexameric rings [3]. DNA may translocate though the 22 Å diameter central pore of this complex. We also solved the crystal structure of TraE, identified the molecular basis of multimerization and described the binding interactions with small molecule inhibitors such as BAR-072 [15, 16]. VirB8 interacts with VirB6 that is the most hydrophobic T4SS component [3]. Due to its hydrophobicity, VirB6 has not been characterized biochemically until now, but according to a long-standing hypothesis in the field it may be part of the DNA transfer pore of the T4SS [3, 14, 17].

We used purified VirB8 and VirB6 homologs TraE and TraD to conduct electrophoretic mobility shift assays (EMSA) and fluorescence polarization (FP) assays to test DNA binding. Both purified proteins bind single-stranded DNA (ssDNA) and double-stranded DNA (dsDNA) in the nanomolar range. We demonstrated that the previously identified T4SS inhibitor BAR-072 inhibits DNA binding to TraE, thus providing insights into its mechanism of action. Finally, we identified conserved amino acids of TraE that are essential for conjugation that may be targets for therapeutic development.

## Results

### TraE binds to ssDNA and dsDNA

TraE was overexpressed and purified in detergents as previously described [3]. To assess if TraE can bind to ssDNA, we carried out EMSA experiments by incubating a constant amount of 5’-fluorescein-labelled DNA (the probe) with increasing amounts of TraE prior to gel migration. To maintain the nucleic acid as single-stranded, we used poly-deoxythymidine oligonucleotides (pT) that have a low propensity to form secondary structure that might impact binding. A 16 nucleotide (nt) pT was selected to ensure the DNA is long enough to span the entire 53 Å length of the hexameric ring as measured by SAXS [3]. We observed slower migration of the 16 nt pT DNA in the presence of TraE in the concentration range of 100–1900 nM (*Figure 1A*). This result suggests that TraE binds to ssDNA, as predicted by the results of previous TrIP experiments [14]. The same experiment was conducted in the presence of dsDNA by annealing the 5’-fluorescein 16 nt pT with 16 nt poly-deoxyadenosine (16 base-pair (bp) pT/pA). Again, we observed slower migration of dsDNA over the same range of TraE concentrations, indicating that TraE also binds dsDNA (*Figure 1B*). TraE appears to bind ssDNA with higher affinity than dsDNA based on the range of concentrations that leads to a complete shift of each probe (compare *Figure 1A to Figure 1B*).

**Legend to Figure 1:**
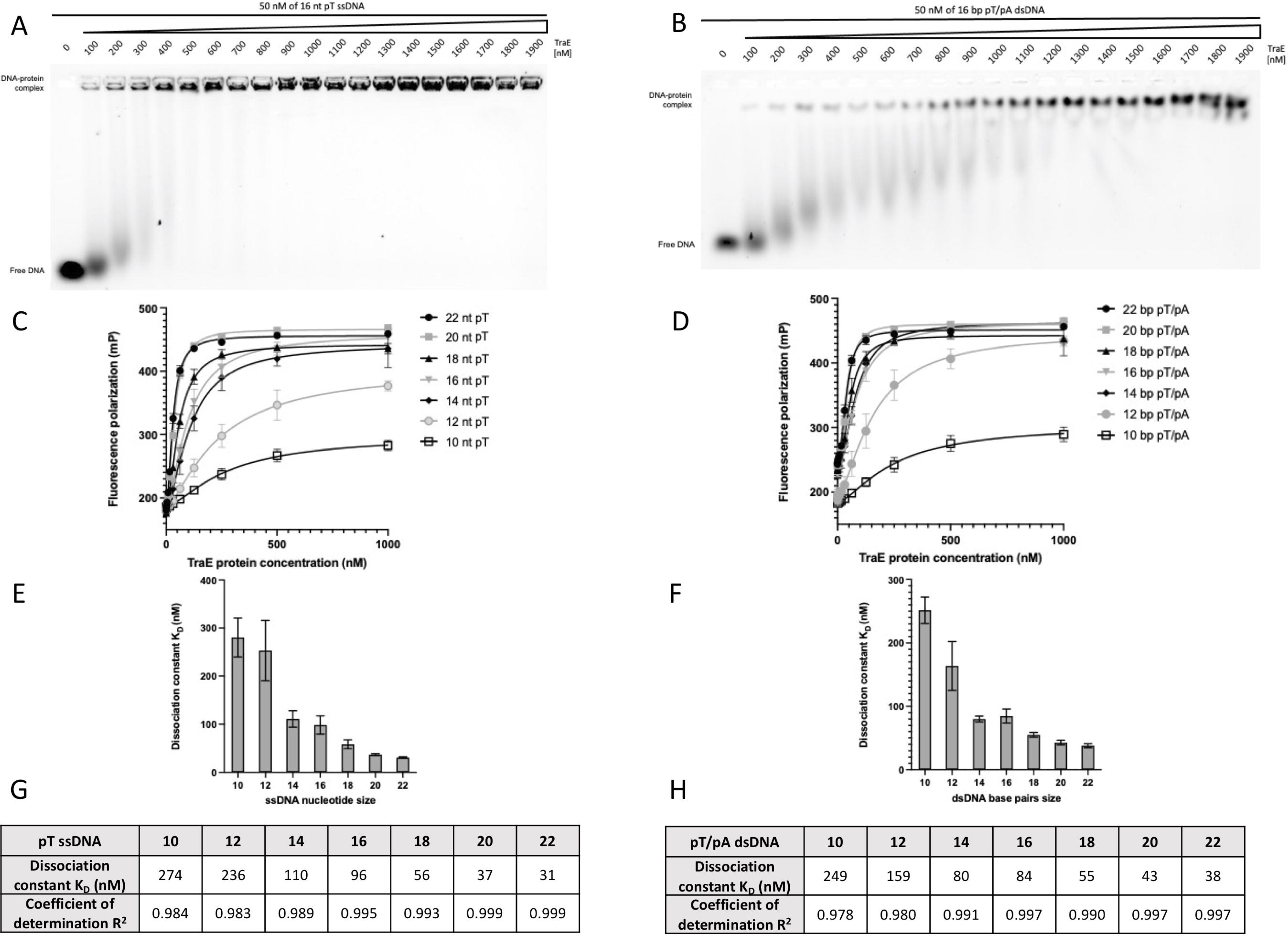
TraE binds ssDNA and dsDNA. EMSA analysis with 50 nM of 16 nt pT ssDNA (**A**) and 50 nM of 16 bp pT/pA dsDNA (**B**) in the presence of TraE gradient from 100–1900 nM. (**C**) FP mP signal of 10–22 nt pT ssDNA and 10–22 bp pT/pA dsDNA (**D**) as a function of TraE protein concentration. Representation of the dissociation constant K_D_ as a function of 10–22 length ssDNA (**E**) or 10–22 length dsDNA (**F**). Summary of TraE K_D_ and R2 of each pT ssDNA (G) and each pT/pA dsDNA (**H**). Each experiment was independently repeated three times, and each value represent the mean of those experiments.

Next, an orthogonal FP DNA-binding assay was conducted to corroborate the EMSA results and to quantify TraE affinity for DNA. According to SAXS data, the central pore has a diameter of 22 Å and a length of 53 Å. Therefore, we decided to use a range of DNA oligomers expected to approximate the length of the central pore, ranging from 10– 22 nt or bp, resulting in theoretical lengths of 34–78 Å. We observed hyperbolic curves for titrations of TraE with ssDNA and dsDNA, which indicated a binding interaction and confirmed our EMSA results. A one-to-one binding model was fit to the data to calculate apparent binding affinities in terms of an equilibrium dissociation constant, K_D_. The binding affinities are in the nanomolar range, largely following an overall DNA length-dependent binding affinity with longer DNA having higher affinity and thus lower K_D_ values. The longest DNAs used (22 nt/bp) resulted in the highest affinities measured: 31 nM for ssDNA and 38 nM for dsDNA (*Figures 1C–H*).

### BAR-072 inhibits TraE DNA binding

In our previous study using the periplasmic version of TraE (TraEp) [15] we discovered that the small molecule BAR-072 binds to TraEp and inhibits *in vivo* bacterial conjugation. We tested whether BAR-072 could inhibit TraE interaction with DNA. For EMSA experiments illustrated in *Figures 2A and 2B*, the concentration of TraE was fixed at 75 nM and then incubated with increasing concentrations of BAR-072. Following incubation and gel migration, we observed more unbound DNA with increasing BAR-072 concentrations, which showed that BAR-072 inhibits TraE-DNA interaction.

**Legend to Figure 2:**
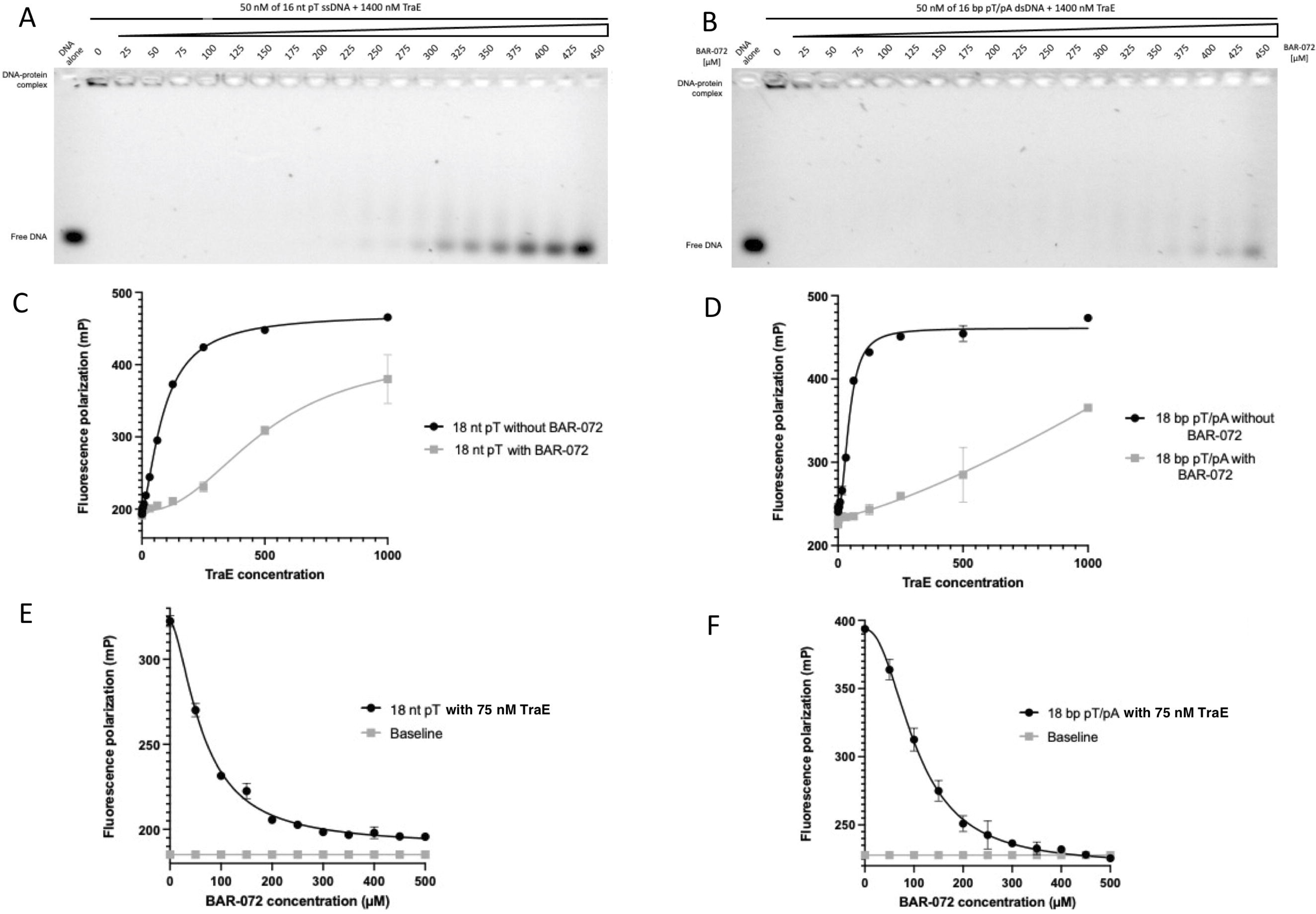
BAR-072 compound inhibits TraE-DNA binding. EMSA analysis of 1400 nM TraE with 50 nM of 16 nt pT ssDNA (**A**) and 50 nM of 16 bp pT dsDNA (**B**) in the presence of BAR-072 gradient ranging from 25–450 μM. FP mP signal of 18 nt pT ssDNA (**C**) or 18 bp pT dsDNA (**D**) as a function of TraE gradient concentration in the presence (grey curve) or absence (black curve) of 450 μM BAR-072 compound. (**E**) FP mP signal of 75 nM TraE incubated with 18 nt pT ssDNA (**E**) or 18 bp pT dsDNA (**F**) as a function of BAR-072 gradient (black curve). The baseline (grey line) represents the total fluorescence emission value in the presence of various BAR-072 concentrations. Each experiment was independently repeated three times, and each value represent the mean of those experiments.

The same experiment was conducted using FP in *Figures 2C and 2D*, but now with 18 nt pT ssDNA and 18 bp pT/pA dsDNA, respectively. A control condition without BAR-072 was included (dark curve in *Figures 2E and F*). In the presence of BAR-072 (grey curves in *Figure 2E and 2F*), we observed a drop in FP signal that correlated with an increasing amount of the compound. The baseline traces in *Figures 2E and 2F* show that BAR-072 does not emit or influence the total fluorescence, as the FP signal does not change over the concentration range tested. These results indicate that BAR-072 inhibits TraE binding to DNA, and we observe that inhibition has a higher impact on binding to ssDNA than on dsDNA.

### TraD binds to ssDNA and dsDNA

Due to the spatial proximity of TraE and TraD in the T4SS, we investigated whether TraD also binds DNA. TraD was overexpressed and purified in various detergents (data not shown) and only Triton X-100 detergent yielded a stable purified protein with a highly homogeneous gel filtration profile (supplementary Fig. 1). As for TraE, an EMSA experiment was first used to evaluate the DNA binding capacity of TraD. In *Figures 3A and 3B*, we observed that TraD retarded DNA migration, indicating that TraD is also capable of binding ssDNA and dsDNA, respectively. FP experiments were also conducted (*Figures 3C and 3D*) to quantify the binding affinity. These experiments showed a binding affinity in the nanomolar range of TraD for both ssDNA and dsDNA (*Figures 3E–H*). Furthermore, as for TraE, TraD has a higher affinity for ssDNA compared to dsDNA for the longest DNA lengths tested.

**Legend to Figure 3:**
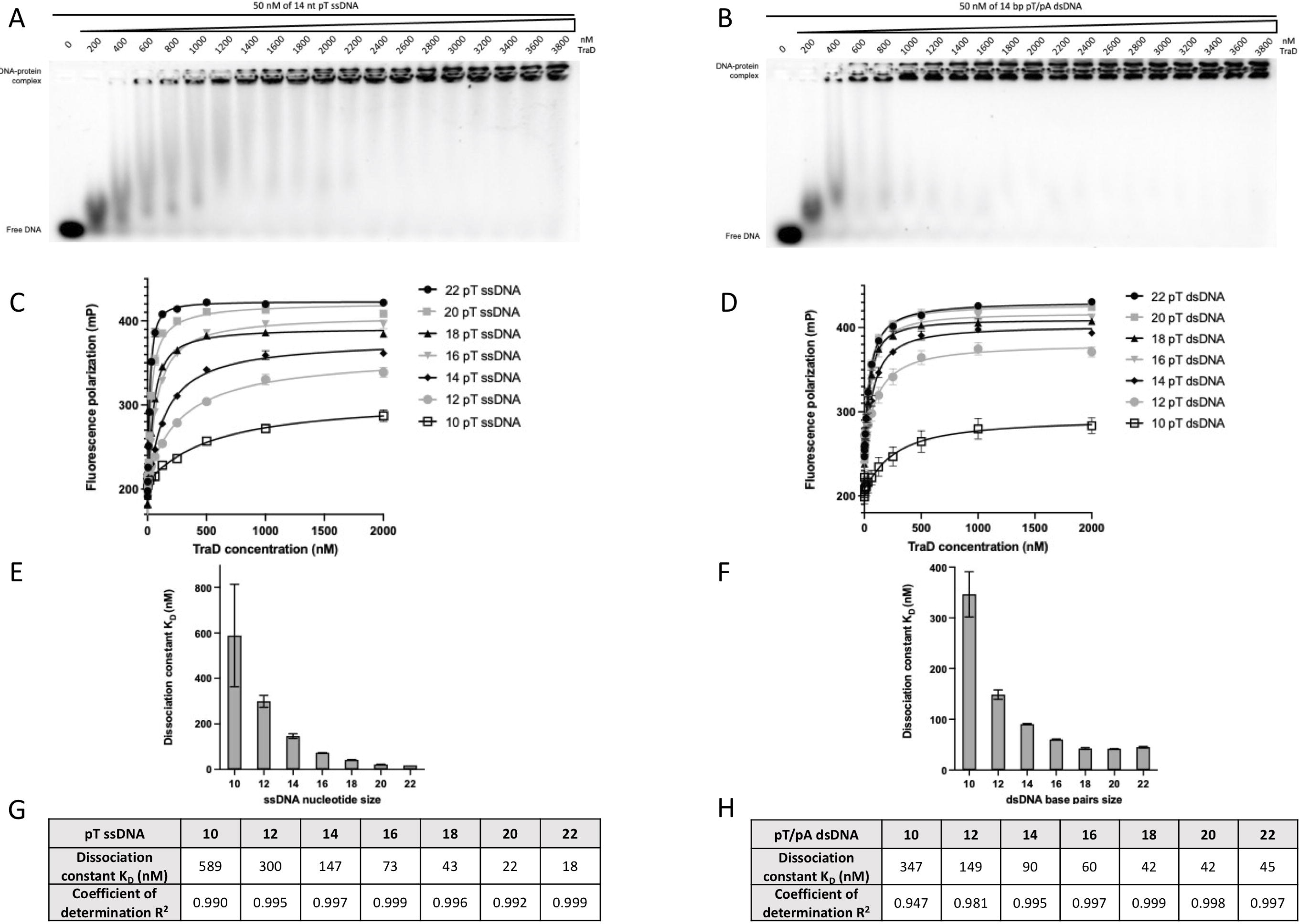
TraD binds ssDNA and dsDNA. EMSA analysis with 50 nM of 14 nt pT ssDNA (**A**) and 50 nM of 14 bp pT dsDNA (**B**) in the presence of TraD gradient from 200–3800 nM. FP mP signal of 10–22 nt pT ssDNA (**C**) and 10–22 bp pT dsDNA (**D**) as a function of TraD protein concentration. Representation of the dissociation constant K_D_ as a function of 10–22 length ssDNA (**E**) or 10–22 length dsDNA (**F**). Summary of TraD K_D_ and R2 of each pT ssDNA (G) and each pT/pA dsDNA (**H**). Each experiment was independently repeated three times, and each value represent the mean of those experiments.

### TraE sequence alignment and mutagenesis

To gain insights into the molecular basis of DNA binding, a multiple sequence alignment of TraE homologs (VirB8 proteins) from different organisms was performed, and several amino acid residues were found to be highly conserved (*Figure 4A*). Since TraE binds DNA non-specifically, we created single point mutations of each positively charged and aromatic conserved amino acid by substituting them with alanine. To determine if these 14 single point mutations impact conjugation, we co-expressed the TraE mutants in a donor strain carrying a Δ*traE* version of pKM101. We incubated the co-transformed bacterial donor cells overnight with recipient cells and tested the efficiency of complementation by plating on media with selective antibiotics to monitor the transfer of the pKM101 Δ*traE* plasmid into the recipients. The efficiency of conjugation was significantly reduced in the case of 13 of 14 TraE mutants (*Figure 4B*) and their localization on the X-ray structure of TraE is shown in *Figure 4C*. Plasmids carrying mutations W40A, Y106A and Y225A complemented conjugation at strongly reduced levels and no complementation was observed in the case of TraE mutants Y113A, R176A, Y203A and R213A. Western blotting with TraE-specific antisera showed that all mutants were expressed at comparable levels (*Figure 4B*), indicating that the mutation do not impact TraE expression. Further work is required to determine if the identified amino acids participate in DNA binding or possibly in another biological role, such as protein-protein interactions.

**Legend to Figure 4:**
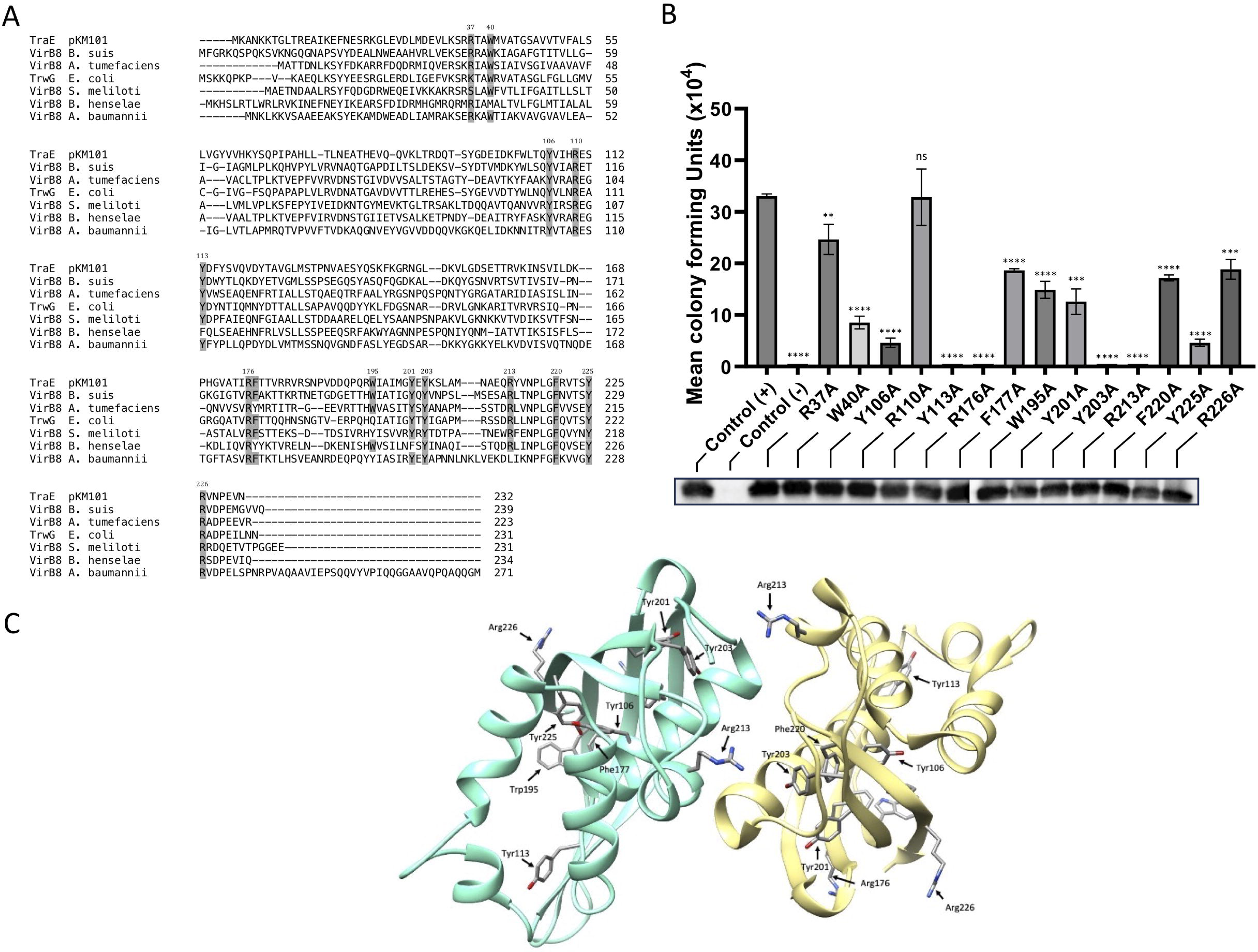
Site-directed mutagenesis on TraE and localization on crystal structure. (**A**) Amino acids sequence alignment of TraE from pKM101 plasmid with six VirB8 homologs from *B. suis*, *A. tumefaciens*, *E. coli* R388 plasmid, *S. meliloti*, *B. henselae*, and *A. baumannii*. Strongly positively charged and aromatic amino acids residues are marked in grey. (**B**) pKM101 conjugation assay to assess the capacity of TraE and its variants to complement a pKM101Δ*traE* plasmid. The data represent averages and standard errors of three biological replicate cultures. Asterisks represent statistically significant differences (**** for p<0.0001, n=3; *** for p<0.001, n=3; ** for p<0.01, n=3; ns for non significant, n=3). Bottom, Western blot using anti-TraE antibody. (**C**) Crystal structure of TraE dimer (pdb: 5I97) with location of mutation sites.

## Discussion

Here, we present direct evidence showing that purified TraE and TraD bind DNA. The binding affinities are in the nanomolar range, suggesting that both proteins are part of the transmembrane substrate translocation channel. Considering the biological role of the relaxosome complex [12], we expected to observe only ssDNA binding. However, a recent preprint article on the outer membrane protein CagX (VirB9 protein homolog in the *Helicobacter pylori* T4SS) shows that CagX can bind ssDNA and dsDNA and the dissociation constants are in the micromolar range [18].

Even if our data demonstrate that TraE and TraD both bind dsDNA, we would have expected a larger difference between the binding affinities for the same size oligonucleotide. Yet, the K_D_ values for ssDNA and dsDNA are quite similar with minor variations observed for oligonucleotides of different lengths. The same behavior (ssDNA and dsDNA binding) was observed for TraE purified with different detergent alternatives (amphipols and nanodiscs (data not shown)). Since TraE and TraD are closely juxtaposed in the T4SS [19] and were shown to interact [3, 14, 17] it would be interesting to test for synergy in the future and to assess whether DNA binding affects the conformation of these proteins.

Whereas we described a hexameric TraE by SEC-MALS and negative stain EM [3], the recently published structure of R388 T4SS shows the presence of 24-mer TrwG (VirB8), the homolog of TraE [20]. However, our work on TraE and TraD was done on the isolated proteins and previous studies show the presence of VirB8 at the cell poles in *A. tumefaciens* and its importance for the recruitment of T4SS components [14, 21]. We conclude that VirB6 and VirB8 may form intermediate oligomeric states during the assembly of the inner membrane T4SS complex. Hexameric TraE may represent an early state in the formation of higher molecular weight complexes in the mature T4SS. Indeed, a similar phenomenon was observed for VirB4, which forms dimers as well as hexamers [22–24].

We have previously shown that the periplasmic domain of TraE forms a dimer that is disrupted in the presence of the inhibitor BAR-072 [15]. The small molecule also hinders bacterial conjugation. Here we show that BAR-072 inhibits DNA binding to TraE, probably by disrupting multimerization and this provides insights into its mechanism of action. Interestingly, BAR-072 interacts with R110, a conserved amino acid that was mutated for the conjugative assay (*Figure 4B*). However, the R110A mutation did not significantly impact conjugation, indicating that multiple mutations would be required to mimic the effect of BAR-072.

Sequence similarities between different VirB8 homologs are generally low and range from 20–40%, but their overall structures are highly conserved suggesting evolutionary pressure to conserve protein folding ([25],[26],[27],[28],[29]). Indeed, mixing purified VirB8 homologs resulted in heterodimers, which is consistent with the close structural similarity among homologs [15]. We performed site-directed mutagenesis of 14 highly conserved positively charged and aromatic amino acids that may be involved in DNA binding. The locations of these residues on the TraE crystal structure are represented in *Figure 4C*. Most mutations are in the NTF2 (nuclear transport factor 2) domain, which is known to be present in proteins with disparate functions, such as small molecule binding and enzymatic activities ([30], [31], [32]). According to the sequence alignment in *Figure 4A*, conserved residues R176 and R226 (TraE numbering) correspond to *Brucella suis* residues R179 (R179^bs^) and R230 (R230^bs^), respectively. It was previously shown that R179^bs^ interacts with VirB10 [33] and mutating R230^bs^ inhibits interaction with VirB4 and also strongly reduces *B. suis* intracellular growth [34]. Our observation of decreased and ablated conjugation with TraE R226A and R176A mutants, respectively, (*Figure 4B*) could be explained by breaking similar contacts between components of the pKM101 T4SS.

Bacterial two-hybrid experiments with *B. suis* VirB8 mutants showed that many impact dimer formation [35]. This is the case for Y110^bs^, Y206^bs^ and Y229^bs^ that correspond to TraE residues Y106, Y203 and Y225, respectively. Expression of the Y106A and Y225A TraE mutants led to strongly reduced conjugation, while the Y203A mutant did not complement at all. These mutations likely reduced TraE multimerization, thereby impacting bacterial conjugation. Finally, the VirB8 residue W198^bs^ was involved in binding to an inhibitor that impacts dimerization without inducing detectable conformational changes [34]. In our study, the corresponding TraE mutant W195A resulted in a 50% loss of conjugation activity (*Figure 4B*).

This work confirms the importance of conserved residues among VirB8 homologs, especially residues in the NTF2-like domain that are involved in protein-protein interactions. These amino acids are potential targets for the development of small-molecule inhibitors of T4SS. Since all mutants expressed at wild type levels (*Figure 4B*), it would be interesting to purify the mutants to assess whether the TraE mutations impact multimerization and/or DNA binding by gel filtration and FP, respectively. Future research will aim to determine full-length protein structures in the presence of DNA to identify the amino acid residues that are directly implicated in DNA binding.

## Materials and Methods

### Strains, Plasmids and DNA manipulation

The strains and plasmids used are described in Table 1. Briefly, the *E. coli* strain XL-1 blue was used as a host for cloning, and BL21(λDE3) pLEMO was used for TraE and TraD overexpression. The Monarch Plasmid Miniprep kit (NEB) was used to isolate DNA plasmids. All mutations were performed using the QuickChange protocol and verified by automated Sanger DNA sequencing.

**Table 1.**
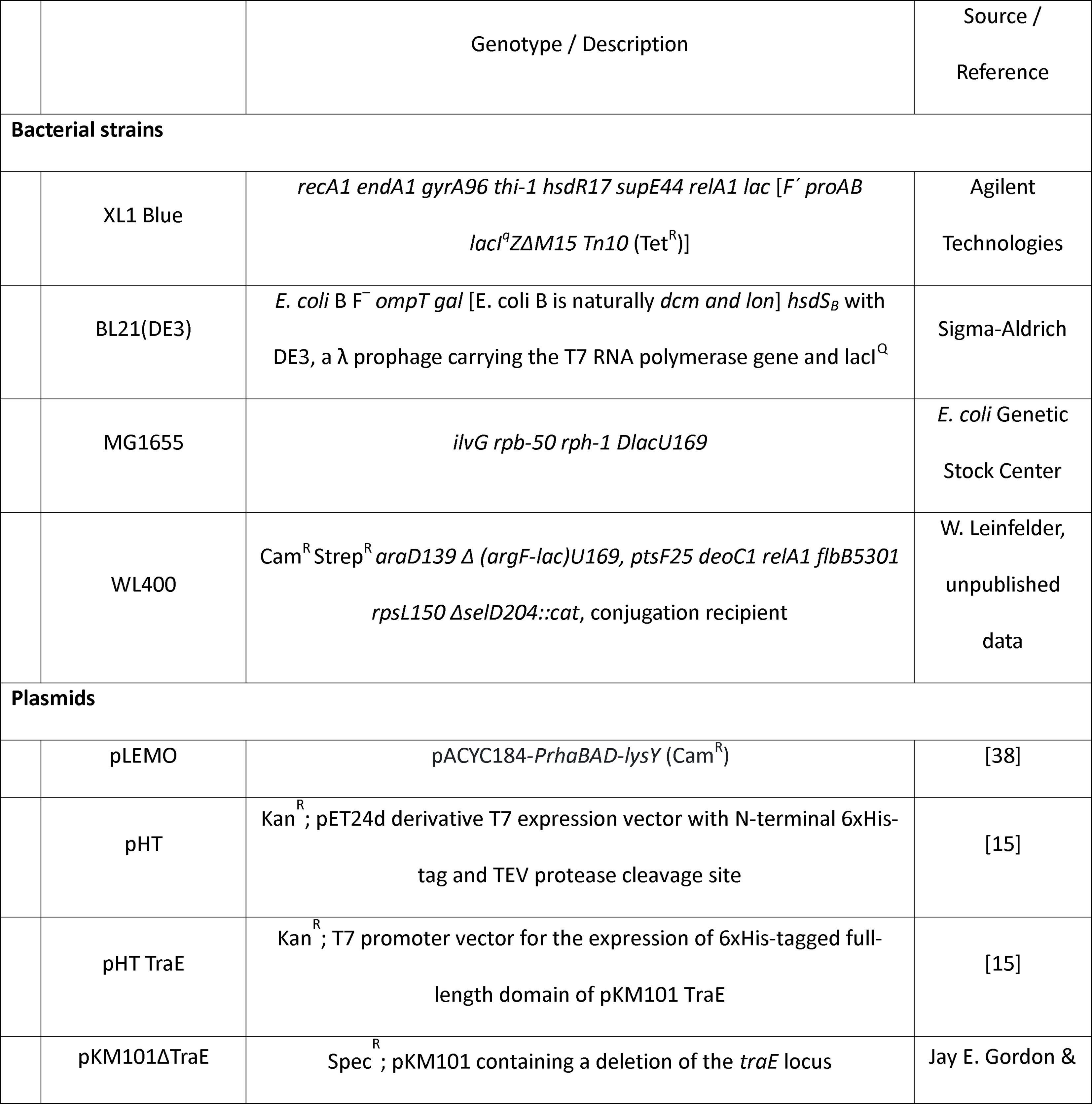
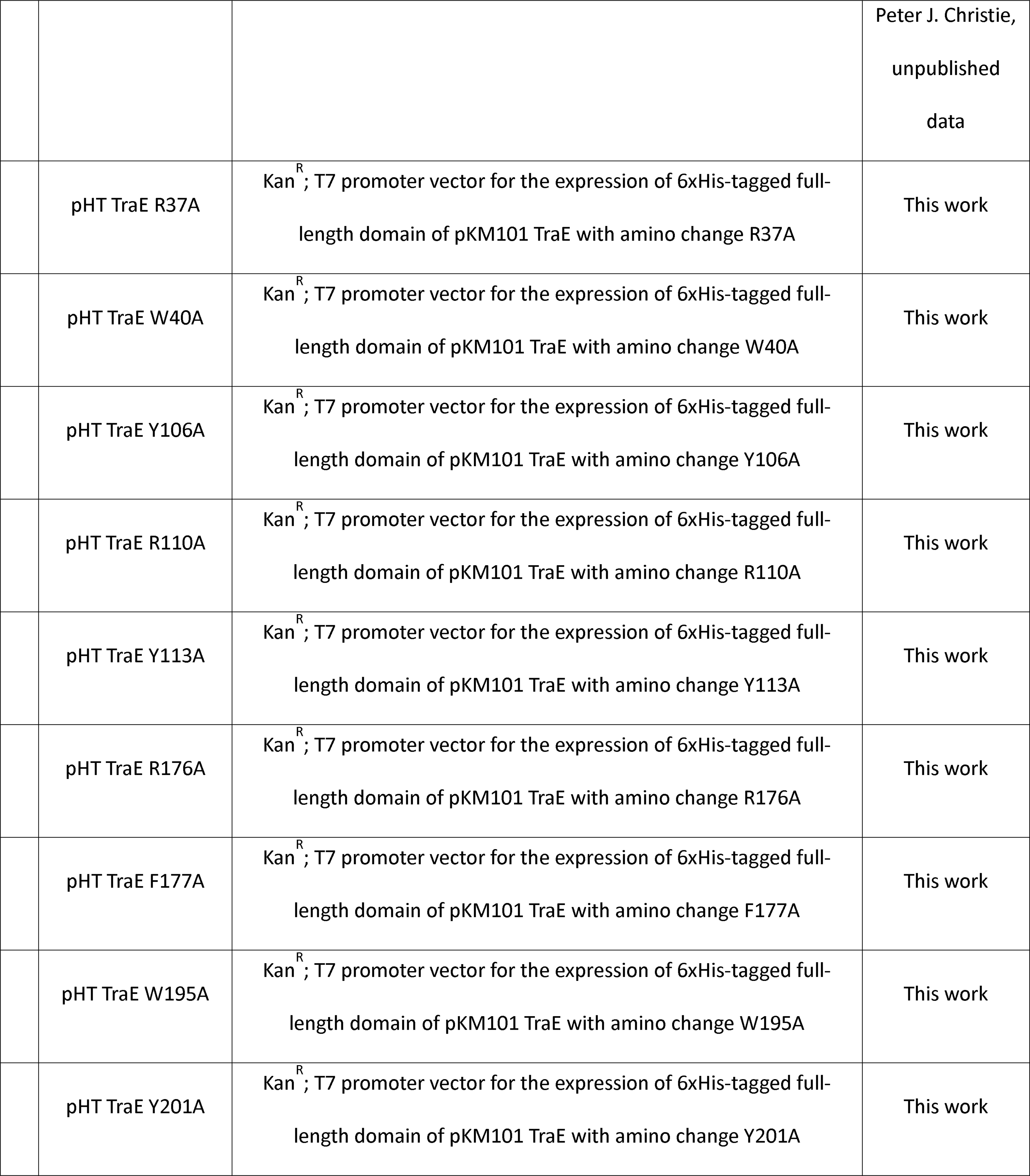
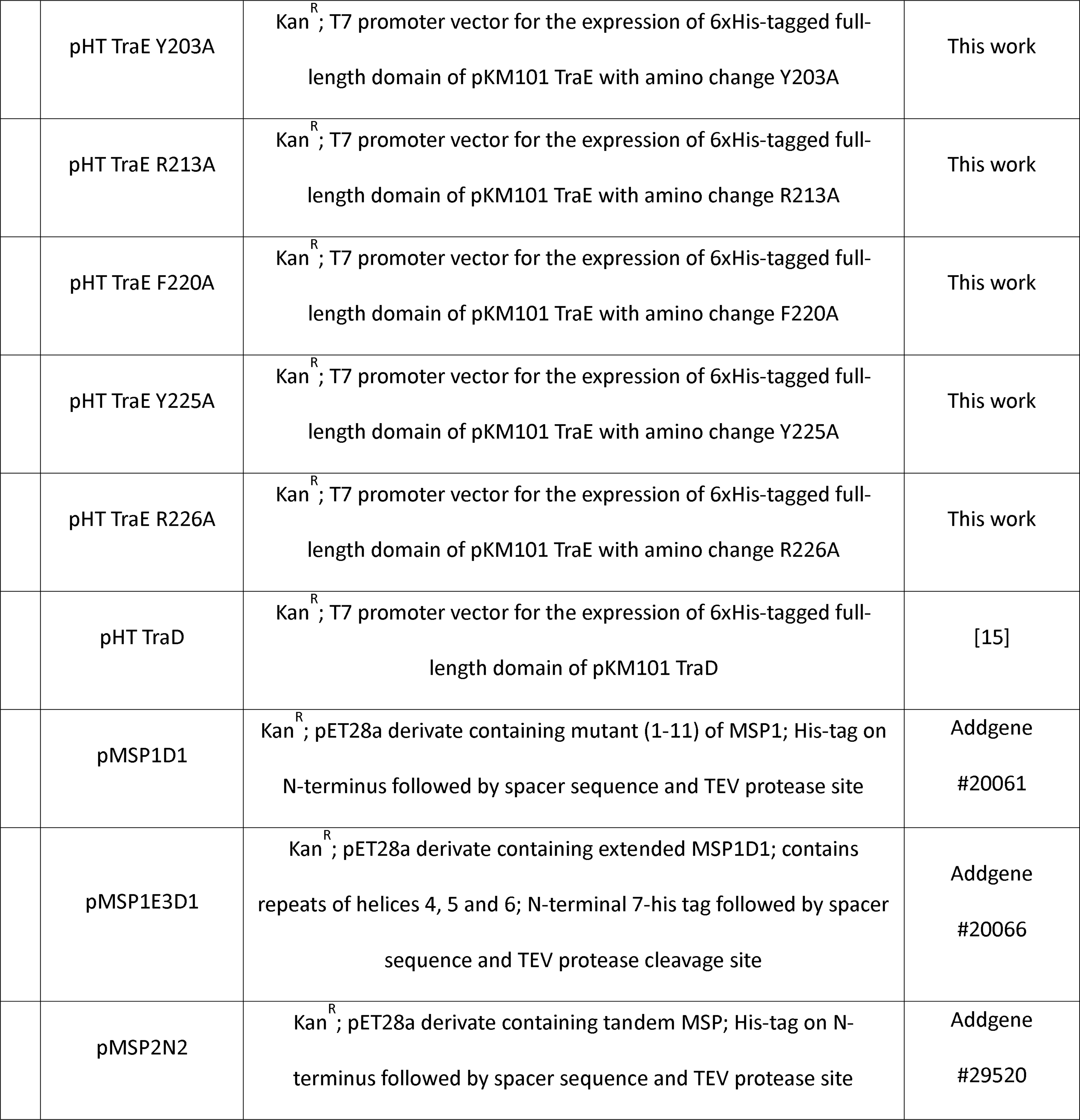
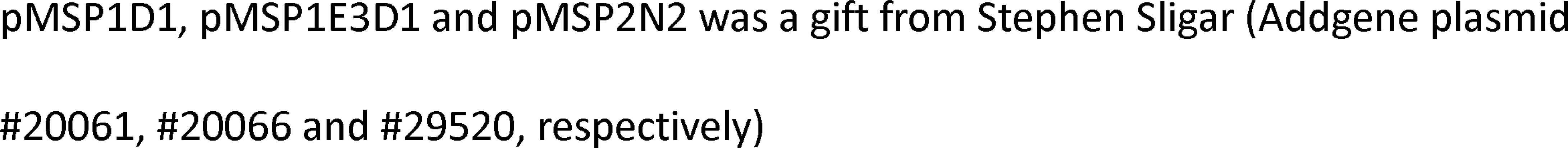
Bacterial strains and plasmids.

### Small-scale TraE and TraD protein expression optimization

TraE and TraD overproduction optimization were performed based on the protocol extensively described by Wang et al., [36]. Briefly, for screening overexpression, the *E. coli* strains BL21(λDE3), BL21 Star(λDE3), C41(λDE3) or C43(λDE3) carrying expression plasmids pHTTraE and pHTTraD were grown under aerobic conditions at 37°C (co-transformed or not with pLEMO plasmid) in 10 mL LB, 2YT or TB media cultures to exponential phase (OD_600_ 0.6–0.8). The expression was induced by the addition of 0, 0.1–1.0 mM IPTG (with 0.1 mM increments); at temperatures of 18°C, 25°C, 30°C or 37°C and cultures were left shaking for 4, 6, 8 or 18 hours at 220 rpm. Cells were collected in 1.5 mL Eppendorf tubes, the DO_600_ was normalized to 1.0 and centrifuged at 16,000 x g for 10 minutes. Pellets were kept at –20°C until further use. Bacterial lysis was performed using the method described by Casu et al., [3]. Dot Blot was performed by pipetting and depositing 3 µL of the bacterial lysis supernatant on nitrocellulose membrane and leaving to air dry for 10 minutes. Blocking and further procedures were the same as for the Western Blot using an anti-His-Tag antibody (1:5000 dilution). Five of the most intense culture conditions visualized by Dot Blot were loaded on SDS-PAGE followed by Western Blot to ensure the protein migrated at the correct molecular weight. For pHTTraE, the best overexpression condition corresponded to the BL21(λDE3) pLEMO induced with 0.5 mM IPTG at 30°C for 6 hours in LB media, while for pHTTraD, the best overexpression condition corresponded to BL21(λDE3) pLEMO induced with 0.5 mM IPTG at 18°C for 18 hours in TB media.

### TraD solubilization tests and purification

Five different detergents (Pluronic F-127, Tergitol NP-40, Triton X-100, DDM, OGNG and LMNG) were used to assess TraD extraction and stabilization. Briefly, bacterial cultures were resuspended at 4°C in Buffer A (50 mM sodium phosphate pH 7.4, 300 mM NaCl, 100 mM imidazole) with cOmplete Mini protease Inhibitor mixture and DNase I at 100 µg/mL and lysed twice using One Shot cell disruptor (Constant Systems, Inc) at 27 kpsi and 4°C. Ultracentrifugation at 250,000 x g for 1 hour at 4°C was performed, and pellets were collected and solubilized overnight at 4°C in Buffer A supplemented with 20x CMC of the detergent. Debris were removed by centrifugation at 34,000 x g for 30 minutes at 4°C. A fraction of the solution was used for Western Blot analysis to assess the quantity of extracted and solubilized TraD, and the rest was loaded on HisTrap Ni-chelate column (GE Healthcare). After extensive washing in Buffer B (Buffer A plus 3x detergent CMC), the protein was eluted into fractions using Buffer C (50 mM sodium phosphate pH 7.4, 300 mM NaCl, 100 mM arginine, 100 mM glutamate, 10% glycerol, 1 M imidazole and 2x detergent CMC). The fraction with the highest concentration was injected into a Superose 6 10/300 column equilibrated with 50 mM HEPES pH 8.0, 150 mM NaCl, 50 mM arginine, 50 mM glutamate, 50 mM imidazole, and 2x detergent CMC.

### TraE purification in OGNG detergent

TraE was purified as described by Casu et al., [3].

### Protein molarity quantification and visualization

All protein molar quantifications were based on the hexameric form of TraE and the pentameric form of TraD (as described in [19]). SDS-PAGE gels were visualized with zinc imidazole [37].

### Electrophoretic mobility shift assay for TraE and TraD

The EMSA was conducted using 50 nM of 5’ 6-Fluorescein (5’ 6-FAM) pT labelled DNA (hybridized or not with the complementary non-fluorescent DNA strain to form ssDNA and dsDNA) from IDT. LMNG was used as a detergent for TraE, while Triton X-100 was used for TraD. Proteins were diluted in 10 mM Tris pH 9.5, 150 mM NaCl, 10% glycerol, 5x detergent CMC and incubated with DNA for 30 minutes at 4°C. Samples were loaded on a 0.5% agarose gel and run in the running buffer: 0.5X Tris-Borate-EDTA (TBE) pH 9.5, 150 mM NaCl, 5% glycerol, 5x detergent CMC at 90V, 4°C for 2 hours. Agarose gels were prepared by solubilizing agarose with the running buffer, and gel revelation was acquired using a Fluorescein filter from the ChemiDoc MP imaging system. For BAR-072 inhibition experiments, all solutions and gels were supplemented with 10% DMSO.

### Fluorescence polarization of TraE and TraD

As for EMSA experiments, 5’ 6-FAM was used but at 10 nM concentration and all fluorescence polarization experiments were performed at least three times. Protein-DNA binding was performed in FP Buffer: 10 mM sodium phosphate (NaPi) pH 7.4, 150 mM NaCl, 10% glycerol and 5x detergent CMC (LMNG for TraE and Triton X-100 for TraD). The probe was incubated with the indicated protein concentration, and measurement was made at room temperature on a Victor3Vplate reader (PerkinElmer). For binding and inhibition, curves were fit with saturation binding, specific binding with Hill slope equations. For inhibition experiments, 10% DMSO was supplemented with the FP buffer.

### Conjugative DNA transfer and Western blot

For conjugative assays, pHT plasmids (kanamycin-resistant) and MG1655 pKM101Δ*traE* (donor, spectinomycin-resistant) were used with WL400 cells (recipient cell, chloramphenicol-resistant). *E. coli* strains MG1655 containing pKM101Δ*traE* were transformed with complementation vector pHT (negative control), pHTTraE (positive control), or pHTTraE expressing TraE variants. The same protocol described by Casu et al., [15] was used, except the recipient cell was grown in liquid LB media containing 50 µg/mL kanamycin and 100 µg/mL spectinomycin and the agar plates for conjugation transfer quantification containing kanamycin and chloramphenicol antibiotics.

For the Western Blot, MG1655 pKM101Δ*traE* transformed with pHT, pHTTraE and pHTTraE variants were grown on LB agar plates (with 100 µg/mL spectinomycin and 50 µg/mL kanamycin) at 30°C overnight. Cells were pelleted, and bacterial lysis was performed [3]. 50 µL of sample was loaded on SDS-PAGE gel and anti-TraE antibody was used (1:5000 dilution).

## Acknowledgments

This work was supported by Canadian Institutes of Health Research grants #274108 and #469471 to (C.B.). pKM101ΔtraE plasmid was kindly provided by Dr. Peter J. Christie (University of Texas Health Science Center at Houston). We thank Antonio Nanci (University of Montreal) and the FEMR staff (McGill University) for advice on single-particle EM analysis.

