## supplementary information for "Inner membrane components of the plasmid pKM101 type IV secretion system TraE and TraD are DNA-binding proteins"

**Supplementary Figure 1**

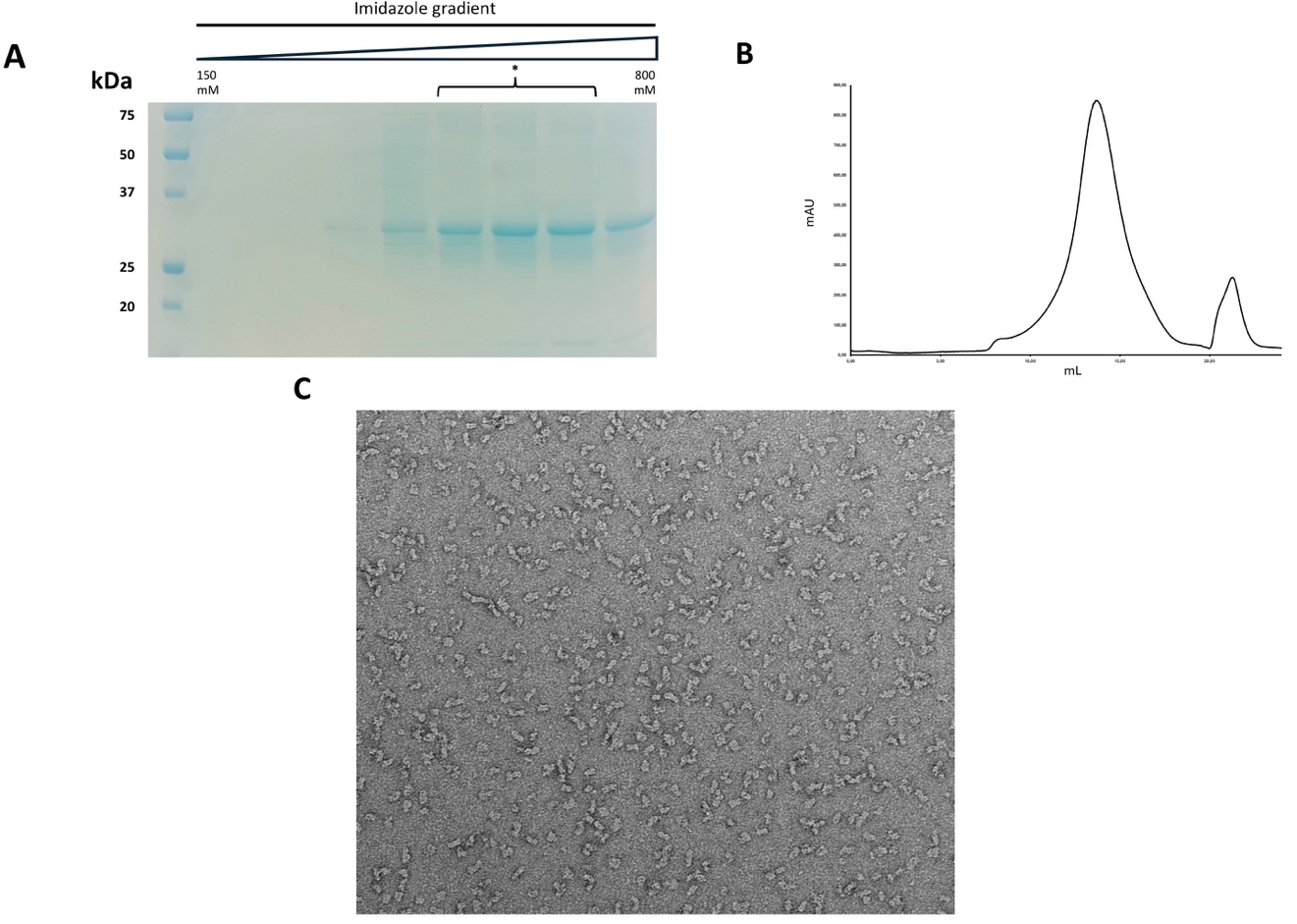

**Legend to supplementary Figure 1:** **TraD purification and analysis.** (A) IMAC elution fractions of TraD analysed on SDS-PAGE with Coomassie blue coloration. (B) Elution profile of TraD on a Superose 6 column. (C) Example of a negative stained micrograph of diluted sample from (B). The (*) labeling represents the fractions collected after elution and injected onto the Superose 6 column.
